# Chronic antidepressant treatments rescue reduced REM sleep theta power in a rat social defeat stress model of depression

**DOI:** 10.1101/2020.03.21.001164

**Authors:** Yoshiki Matsuda, Nobuyuki Ozawa, Takiko Shinozaki, Kazuhisa Aoki, Naomi Nihonmatsu-Kikuchi, Toshikazu Shinba, Yoshitaka Tatebayashi

**Affiliations:** Affective Disorders Research Project, Tokyo Metropolitan Institute of Medical Science, 2-1-6 Kamikitazawa, Setagaya, Tokyo, 156-8605, JAPAN; Department of Psychiatry, Shizuoka Saiseikai General Hospital, 1-1-1 Oshika, Suruga-ku, Shizuoka, 422-8527, JAPAN

## Abstract

Although It is widely recognized that virtually all antidepressants (ADs) suppress rapid eye movement (REM) sleep, the effects of chronic AD treatments on sleep abnormalities, especially those of REM sleep, have rarely been investigated comprehensively in animal models of major depressive disorder (MDD). Here, we show that chronic social defeat stress (SDS) in rats induces prolonged social avoidance and MDD-like alterations in sleep architecture (increased REM sleep durations, bouts, and shortened REM latency) even a month after the last SDS. These abnormalities were associated with changes in electroencephalography (EEG) spectra powers, such as reduced REM sleep theta powers during the light phase. Chronic AD treatments significantly ameliorated these behavioral, sleep, and EEG abnormalities, although in some cases not to control levels. Interestingly, the social interaction ratios a month after the last SDS most strongly correlated with the REM sleep theta powers. These results suggest that chronic AD treatments suppress REM sleep durations and bouts as observed in human MDD patients, but, at the same time, increase REM sleep theta power. The latter is an EEG parameter that has never been directly investigated in humans, and which may be responsible for the therapeutic effects of ADs.

**Significance Statement:** Disturbed sleep is one of nine diagnostic criteria for major depressive disorder (MDD), which usually resolves with adequate treatment. However, little is known about how antidepressants (AD) work in MDD animal models. Here, we developed a novel rat social defeat stress model demonstrating long-lasting MDD-like sleep disturbances. We found that chronic AD treatments suppressed REM durations and bouts as reported previously, but, at the same time, increased REM sleep theta power. Interestingly, among several sleep parameters, only REM sleep theta power strongly correlated with depressive symptoms at 1M. Thus, REM sleep theta power, an EEG parameter that has never been investigated directly in humans, could be a novel indicator for MDD and/or AD effects.

## Introduction

Major depressive disorder (MDD) is common, with a 12-month prevalence as high as 3.2% in subjects without a comorbid physical disease, and between 9.3 and 23.0% in those with such conditions [Moussavi et al., 2007]. Despite its frequency, treatment options for MDD remain unsatisfactory for many patients due to the high likelihood of a chronic course, negative impacts on quality of life, and heightened suicide risk. Disturbed sleep is reported by up to 90% of MDD patients [Riemann et al.,2001; Tsuno et al., 2005] and increases risks of the development [Ford and Kamerow, 1989] and relapse of MDD [Peterson and Benca, 2008].

The most common electroencephalogram (EEG) sleep abnormalities in MDD include decreased rapid eye movement (REM) sleep latency (= interval between sleep onset and the first REM sleep period [Kupfer et al., 1982]), increased total REM sleep time and REM density (i.e., frequency of rapid eye movements per REM period), and decreased non-REM (NREM) sleep [Benca et al., 1992; Benca, 2005; Peterson and Benca, 2008]. These changes may be predictive of poorer clinical outcomes for psychotherapy [Thase et al., 1997] and their normalization may be an early predictor of antidepressant drug (AD) response [Krystal et al., 2008]. Recent meta-analyses, however, indicated that no single sleep variable reliably distinguishes patients with MDD from healthy controls [Pillai et al., 2011; Baglioni et al., 2016].

Virtually all ADs suppress REM sleep (duration) [Tsuno et al., 2005; Palagini et al., 2013]. This effect, likely mediated by serotonin [5-hydroxytryptamine (5-HT)] and/or noradrenalin [McGinty and Harper, 1976; Jones, 1991; Thakkar et al., 1998; Landolt et al., 2003], is rapid [Mayers and Baldwin, 2005] and continuous for at least 4 – 8 weeks [Stassen et al., 1993; Beitinger and Fulda, 2010]. In the CNS, the raphe, a diffuse network of brainstem nuclei, synthesizes 5-HT and innervates and receives reciprocal inputs from many brain regions [Azmitia and Siegel, 1978; Weissbourd et al., 2014]. The 5-HT system is wake-active [Sakai, 2011] and forms part of the ascending arousal system promoting wakefulness [Scammell et al., 2017]. A recent study, however, demonstrated that the 5-HT system, especially its tonic stimulation, promotes sleep by generating homeostatic sleep pressure during wakefulness [Oikonomou et al., 2019]. Thus, the actual roles of 5-HT on sleep, particularly those elicited by chronic AD treatments, remain elusive.

To untangle these multifaceted relationships in MDD, valid animal models will be critical. A rodent social defeat stress (SDS) model may prove the most effective because, unlike all other rodent chronic stress models, in which behavioral abnormalities rapidly revert to normal days after the last stress, a subset of behavioral abnormalities induced by SDS remains nearly permanent, which can be reversed by chronic AD treatments [Berton et al., 2006; Tsankova et al., 2006; Krishnan et al., 2007; Wilkinson et al., 2009; Vialou et al., 2010; Cao et al., 2010]. However, little is known about the post-stress effects of SDS on sleep and/or circadian rhythm. For example, in mouse SDS models, dysregulation in the circadian amplitude of body temperatures was noted, but disappeared naturally 4 weeks after the last SDS [Krishnan et al., 2007]. In rat models, effects like reduced circadian amplitudes in home cage activity, body temperature and heart rate, can last for a few weeks after the last SDS [Tornatzky and Miczek, 1993; Harper et al., 1996; Meerlo et al., 1997; Meerlo et al., 2002]. While these circadian changes may be associated with alterations in sleep/wake states, no study has examined EEG sleep alterations as well as the effects of chronic AD treatments in these models.

Here, we established a novel rat SDS model revealing long-term post-stress social avoidance linked to MDD-like sleep disturbances, investigated the effects of chronic AD treatments, and found that, among several sleep-related variables, decreased REM sleep theta power, a parameter never investigated directly in human MDD patients, may be a core sleep symptom contributing to MDD-like behaviors in rats.

## Methods and Materials

### Animal care and use

All animals were treated according to the protocols and guidelines approved by the Animal Use and Care Committee of the Tokyo Metropolitan Institute of Medical Science. All animals were kept under standard laboratory conditions (12 h light/dark (LD) cycle (LD12:12); lights on at 8:00 a.m. (= Zeitgeber time 0 (ZT0))) with food and water available ad libitum unless otherwise indicated.

### Social defeat stress

A 14-day repeated social defeat stress (SDS) paradigm, originally developed for mice [Berton et al., 2006; Krishnan et al., 2007], was applied to rats. Briefly, each male Sprague Dawley (SD) rat (8 weeks old at the onset of stress) was transferred into the home cage of a retired aggressive male Brown Norway (BN) rat (>7 months old; all rats from Charles River Laboratories Japan, Inc., Yokohama, Japan) for 10 min. During this direct contact period, submissive behaviors - including fleeing, crouching and upright posture - were generally observed in the intruder SD rat, signifying it had assumed a subordinate position. If the resident BN rat did not initiate an attack within 5 min, or if the intruder SD rat did not exhibit any submissive behaviors, the SD rat was transferred to a new resident home cage. Other important aspects of the direct contact period were the avoidance of severe injuries and damage to the EEG/EMG electrodes. After 10 min of direct contact, the resident and intruder rats were then kept in indirect contact for 24 hrs using a perforated clear vinyl chloride divider in the resident cage. The next day the intruder SD rat was exposed to a novel resident BN aggressor. Subsequently, intruders were subjected to combined stress (direct and indirect contact) for the first 5 days, followed by only indirect contact for the next two days. This process was continuously repeated for 2 weeks (Fig. 1*A*, S1*A*). Control animals were housed on one side of a perforated divider without a resident. All rats were housed individually except for the SDS period.

**Figure 1.**
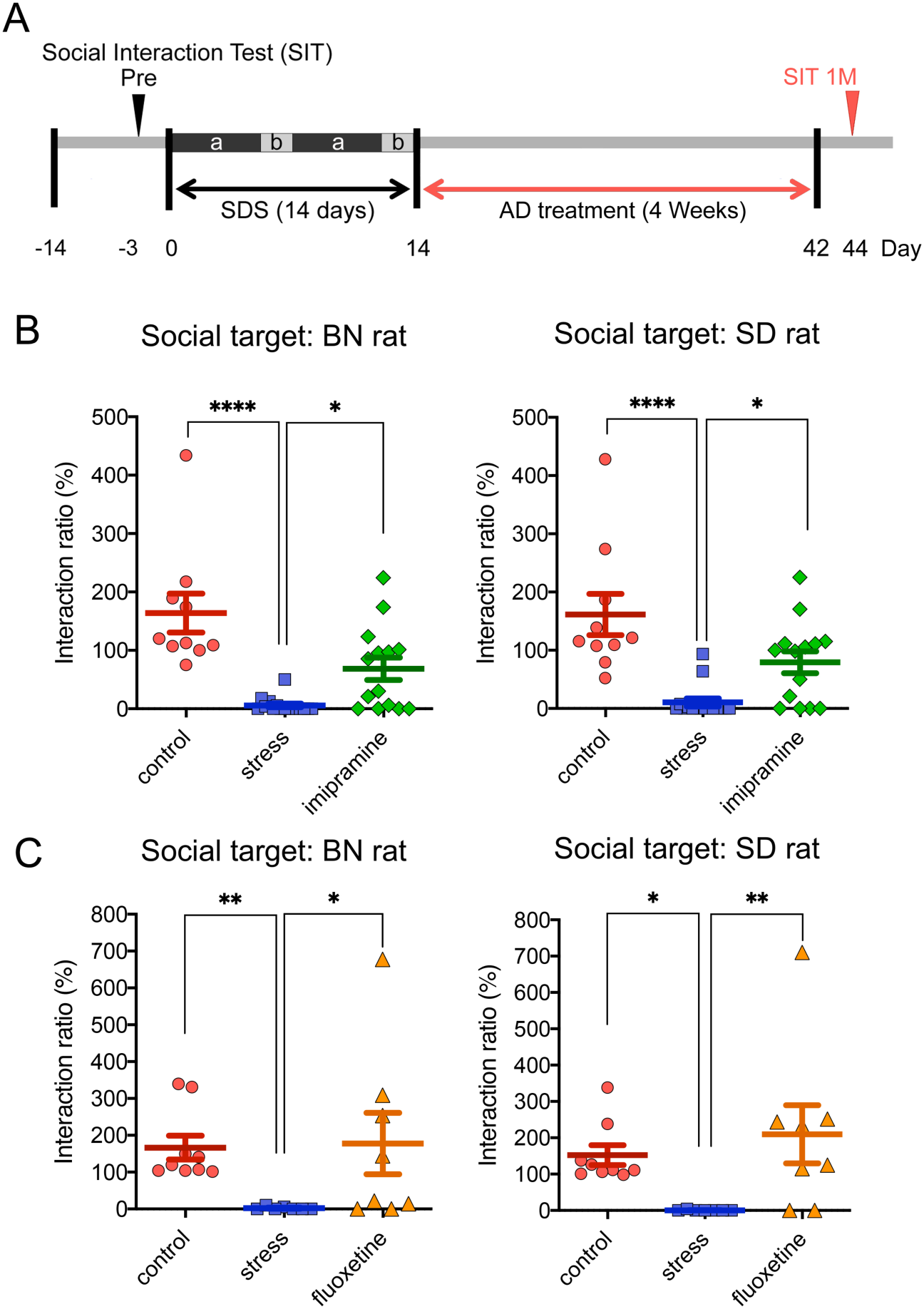
Effects of SDS and chronic AD treatment on behaviors. ***A***, Experimental schedule for behavioral analyses. The 14-day SDS paradigm consisted of 24-h indirect interaction with (a) or without (b) daily 10 min direct interactions for 2 weeks. ***B, C***, Social interaction test one month after the last SDS (Day 44, n = 7 – 16/group). The 14-day SDS induced almost complete social avoidance, whether against an unfamiliar BN (resident; ***B***) or SD (intruder; ***C***) rat, while chronic (4 weeks) IMI or FLU treatment significantly ameliorated this behavior. Note that the AD groups included certain proportions of non-responders. \*\*\**p* < 0.005, \*\**p* < 0.01, \**p* < 0.05; Kruskal-Wallis test followed by Dunn’s multiple comparisons test.

### Chronic AD treatment

After the 14-day SDS paradigm, a tricyclic antidepressant, imipramine (IMI; 20 mg/kg per day (ICN, Aurora, OH, USA)) [Berton et al, 2006], a selective serotonin reuptake inhibitor, fluoxetine (FLU; 5 mg/Kg per day (Cayman, MI, USA)) [Der-Avakian et al., 2014], or saline was intraperitoneally administered for 4 weeks to the SDS rats. Saline was intraperitoneally administered to control rats for 4 weeks.

### Social interactions

An experimental SD rat was placed inside an open field (90 x 90 x 45 cm, 30 lux) and movement was monitored for two consecutive sessions of 2.5 min each. During the first session (“No target”), an empty wire mesh cage (30 x 15 x 15 cm) was placed at one end of the field (Fig. S1*B*). During the second session (“target”), an unfamiliar BN or SD rat was placed in the mesh cage (Fig. S1*B*). Locomotor activity measurements (distance traveled) and time spent in the interaction zone were quantified using custom applications (IMAGE EP and IMAGE FZ, O’hara & co., Ltd., Tokyo, Japan) developed from the public-domain NIH IMAGE program (NIH, Bethesda, USA). The social interaction ratio (%) is defined by the following formula: time (sec) spent in the interaction zone (IZ) with social target (ST) / time (sec) spent in the interaction zone (IZ) without ST x 100 (Fig. S1*B*).

### Surgery and electroencephalogram (EEG) recording

Electrode implantation was conducted as described previously [Shinba, 2009]. Briefly, under pentobarbital sodium (Somunopentyl; Kyoritsu, Tokyo, Japan) anesthesia (60mg/kg, i.m.), rats were fixed to a stereotaxic apparatus (SR-6M; Narishige, Tokyo, Japan). Non-polarized Ag/AgCl screw electrodes (1mm in diameter, O’hara & co., Ltd., Tokyo, Japan) were then implanted epidurally on the left side of the parietal cortex (Br −2.0, L1.5) (Fig. S2*A*). Reference and ground electrodes were placed above the cerebellum (Fig. S2*A*). An electromyography (EMG) stainless electrode surface was subcutaneously placed on the dorsal neck muscle (Fig. S2*A*). Lead wires from all electrodes were soldered to a small square socket and mounted on the skull with acrylic resin cement, along with the electrodes. A recovery period (more than 10 days) was scheduled prior to the start of the experiment.

To collect the EEG data, the rat was moved to a cylindrical experimental cage (35 cm diameter) with a soundproof box kept in LD12:12 (Fig. S2*B*), and connected via a hand-made recording cable with a built-in operational amplifier (TL074; Texas Instruments, Dallas, TX, Fig. S2*B*) in order to lower cable impedance and reduce electrical and movement artifacts. This stage was recorded for 27 hrs (3 hrs habituation and 24 hrs recording), which started at 13:30 and ended at 16:30 (+ 1 day)). Signals were amplified and filtered (gain of 2000 and time constant of 0.1 s for EEG; gain of 5000 and time constant of 0.003 s for EMG; high cut-off filter for both; AB-621G, Nihon-Koden, Tokyo, Japan), digitized at a sampling rate of 500 Hz (Power 1401; Cambridge Electronic Design, Cambridge, UK) and stored using data acquisition software Spike2 (Cambridge Electronic Design, Cambridge, UK). The 27-hr EEG measurements were allocated randomly during days 32 to 42 to match the averaged post-stress days among the groups.

### Sleep scoring

Sleep/wake stages were scored using the automatic scoring protocol incorporated in Spike2 software (rat sleep auto script). These scores were calculated by analyzing the EEG and EMG signals during consecutive 30-second epochs. In addition, the accuracy of the scoring was confirmed manually (Fig. S2*C, D*).

### REM EEG power spectrum

EEG power spectra (0 – 250 Hz) were calculated by using a fast Fourier transform (FFT) function with a Hamming window in the Spike2 software. A block size of 2048 was chosen to allow for a frequency resolution of 0.244 Hz. The Spike2 software script (rat sleep auto script) allowed us to further extract the power spectra per one epoch (30 s) for each sleep/wake stage (NREM, REM, or WAKE). In order to adjust the individual variations, relative power spectra per one epoch were further calculated as the ratio of each power spectra value to the sum of the powers of the band between 2.4 – 30 Hz. The resulting relative power values were averaged for the total REM sleep epochs during the light phase (ZT0 – ZT8.5) and used for the analyses.

### Fear conditioning

On the first day of testing, animals were placed in a sound-attenuated clear acrylic chamber with a grid floor with house lights (20 lux) on and background noise provided by the built-in fan (context A) for 5 min. After 2 min of habituation, rats experienced 2 consecutive tones (white noise, 60 db) for 30 sec, followed by a foot shock of 0.3 mA for 2 sec. The interval between the first and second tone was 30 sec. On the second day, the animals were tested for contextual memory in the same box (context A) for 5 min, but without the tone stimuli. On the third day, the animals were tested for cued memory in the box, now bearing a different appearance (black hollow triangle pole) (context B), but with the tone sounding for a 5-min period. Locomotor activity and freezing behavior were monitored and analyzed as described above.

### Statistical analyses

All data represent the mean ± SEM. All statistical analyses were performed using GraphPad Prism 8 (La Jolla, CA, USA). For statistical comparisons of two groups, the Mann–Whitney U test, as the nonparametric version of the parametric t-test, was used. For datasets that compared more than two groups, the Kruskal-Wallis test, Brown-Forsythe and Welch’s ANOVA test, one- or two-way ANOVA with *post hoc* comparisons were used. The value of *p* < 0.05 was accepted as statistically significant.

## Results

### Establishment of a rat SDS model

To explore effective intruder-resident combinations for the rat SDS paradigm, we first examined several combinations of rats (e.g., Sprague Dawley (SD) vs. SD, SD vs. Wistar, SD vs. Brown Norway (BN)) for a direct interaction period of 10 min. We found that a combination of 10-week-old male SD rats, which served as intruders, and 8-to 12-month-old male BN sexually experienced retired breeder rats, which served as residents, exhibited the stable submissive posture of the SD (test) rats due to attacks by the BN (aggressor) rats. Thus, we decided to apply this combination to a 14-Day SDS paradigm [Berton et al., 2006; Krishnan et al., 2007].

To measure the impact of SDS, the social interaction test [Berton et al., 2006; Krishnan et al., 2007] was conducted just one day after the last SDS. SD (test) rats exhibited near complete social avoidance of unfamiliar resident (BN) rats (After: *p* < 0.0001, Mann-Whitney U test; Fig. S1*C*). Furthermore, this nearly complete social avoidance was also observed with an unfamiliar SD rat (After: *p* < 0.0001, Mann-Whitney U test; Fig. S1*D*), a more universal MDD-like behavioral change. Social avoidance to the BN rat persisted for at least 3 months (1W: *p* < 0.0001; 1M: *p* < 0.0001; 3M: *p* < 0.0001, Mann-Whitney U test; Fig. S1*C*), while that to the SD rat continued for at least a month (1W: *p* < 0.0001; 1M: *p* < 0.0001, Mann-Whitney U test; Fig. S1*D*), although a trend still became apparent at 3 months (3M: *p* = 0.0535, Mann-Whitney U test; Fig. S1*D*). Chronic IMI (Fig. 1*B*) or FLU (Fig. 1*C*) administration for 4 weeks significantly ameliorated the social avoidance, whether to an unfamiliar resident or intruder (stress vs IMI: BN rat: *p* = 0.042, SD rat: *p* = 0.019; stress vs FLU: BN rat: *p* = 0.048, SD rat: *p* = 0.009, Kruskal-Wallis test; Fig. 1*B, C*), although about 30 to 40% showed no improvement (Fig. 1*B, C*).

### Sleep architecture analyses

The sleep/wake stage was determined by using EEG and skeletal muscle atonia in EMG (Fig. S2*A*-*D*). We conducted EEG measurements during 18 to 28 post-stress days (Experimental Day 32 – 42 in Fig. 2*A*) and found that SDS significantly decreased NREM sleep duration (NREM: control (n =5) vs stress (n =6): 5.44 ± 0.12 vs 4.85 ± 0.03, *p* < 0.0001; *F*_(6, 32)_ = 12.20, *p* = 0.0216, two-way (group x sleep stage) ANOVA with Tukey’s multiple-comparisons test; Fig. 2*B*), increased REM sleep duration (REM: control (n = 5) vs stress (n =6): 0.41 ± 0.11 vs 1.49 ± 0.13, *p* = 0.0006; *F*_(6, 32)_ = 12.20, *p* < 0.0001, two-way (group x sleep stage) repeated ANOVA with Tukey’s multiple-comparisons test; Fig. 2*B*) and bouts (REM: control (n =5) vs stress (n =6): 17.20 ± 3.40 vs 57.67 ± 4.82, *p* < 0.0001; *F*_(6, 48)_ = 10.06, *p* < 0.0001, two-way ANOVA with Tukey’s multiple-comparisons test; Fig. 2*C*) during the light phase (ZT0 – ZT8.5). We also found that SDS significantly shortened latency to the onset of the first REM period (control (n =5) vs stress (n =6): 4014.00 ± 630.60 vs 1482.50 ± 375.43, *p* = 0.0259; *p* = 0.0092, Kruskal-Wallis test with Dunn’s multiple-comparisons test; Fig. 2*D*) (For the definition of REM sleep latency, see Fig. S2*D*).

**Figure 2.**
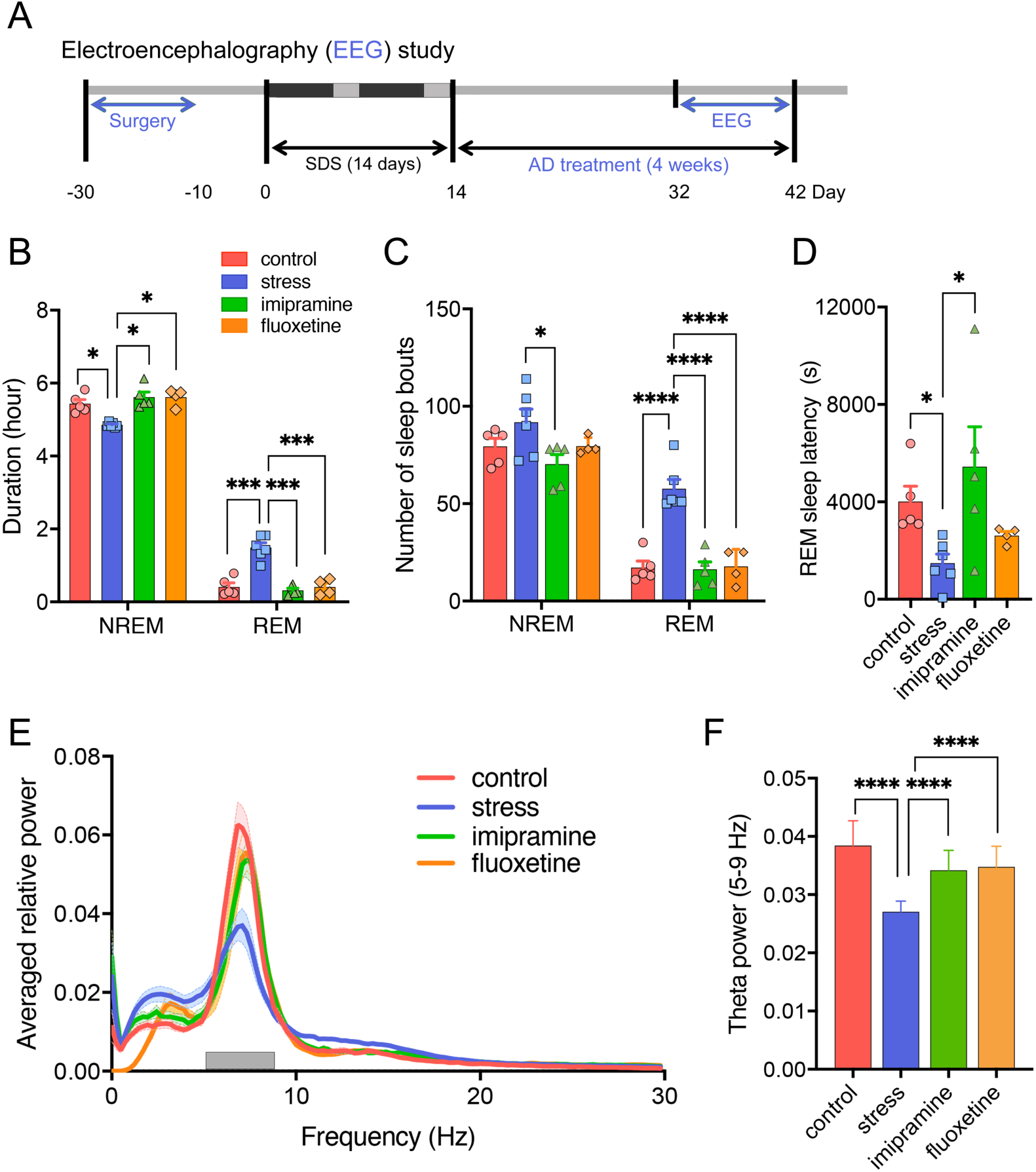
Effects of SDS and chronic AD treatment on sleep architectures and EEG spectral power. ***A***, Experimental schedules for sleep analyses. ***B***, Time spent in NREM, and REM sleep during the light phase (8:00 (ZT0) – 16:30 (ZT8.5)) one month after the last SDS. \**p* < 0.05, \*\*\*\**p* < 0.001, two-way ANOVA followed by Tukey’s test. n = 4 – 6/group. ZT; zeitgeber time. ***C***, Number of sleep bouts in NREM, and REM sleep during the light phase (ZT0 – ZT8.5) one month after the last SDS. \*\**p* < 0.01, \*\*\**p* < 0.005, two-way (number of sleep bouts x group) ANOVA followed by Tukey’s test. n = 4 – 6/group. ***D***, REM latency by SDS and chronic IMI or FLU treatment. \**p* < 0.05, one-way ANOVA followed by Tukey’s test. n = 4 – 6/group. ***E***, Effects of SDS and chronic AD treatments on spectral powers. EEG power spectra during REM sleep in the light phase (ZT0 – ZT8.5) is shown. Gray bars indicate selected theta (5 – 9 Hz) wave lengths. ***F***, Averaged relative spectral power of the theta (5 – 9 Hz) band during REM sleep. Effects of SDS and chronic IMI or FLU treatment. \*\*\*\**p* < 0.001, two-way (Hz x group) ANOVA followed by Tukey’s test.

Chronic AD treatments largely restored most of these sleep alterations to a significant degree (REM sleep duration: stress (n =6) vs IMI (n =5): 1.49 ± 0.13 vs 0.32 ± 0.06, *p* = 0.0003; stress (n =6) vs FLU (n =4): 1.49 ± 0.13 vs 0.41 ± 0.11, *p* = 0.001; *F*_(6, 32)_ = 12.20, *p* < 0.0001, two-way (group x sleep stage) ANOVA with Tukey’s multiple-comparisons test; Fig. 2*B*; number of REM sleep bouts: stress (n =6) vs IMI (n =5): 57.67 ± 4.82 vs 16.20 ± 3.87, *p* < 0.0001; stress (n =6) vs FLU (n =4): 57.67 ± 4.82 vs 17.75 ± 4.42, *p* = 0.0001; *F*_(6, 48)_ = 10.06, *p* < 0.0001, two-way ANOVA with Tukey’s multiple-comparisons test; Fig. 2*C*; REM sleep latency: stress (n =6) vs IMI (n =5): 1482.50 ± 375.43 vs 5442.00 ± 1642.17, *p* = 0.0283; *p* = 0.0092, Kruskal-Wallis test with Dunn’s multiple-comparisons test; Fig. 2*D*), though the restoration in REM sleep latency proved much less significant (e.g. FLU: stress (n =6) vs FLU (n =4): 1482.50 ± 375.43 vs 2617.50 ± 163.11, *p* > 0.999; *p* = 0.0092, Kruskal-Wallis test with Dunn’s multiple-comparisons test; Fig. 2*D*). We also investigated time-dependent effects of chronic FLU treatments on EEG sleep parameters (Fig. S3*A*) and found that FLU restored NREM duration slowly (after stress (n = 4) vs 1W FLU (n = 4): 4.92 ± 0.31 vs 5.05 ± 0.09, *p* = 0.9613; 1W FLU (n = 4) vs 1M FLU (n = 4): 5.05 ± 0.09 vs 5.62 ± 0.12, *p* = 0.0237; *p* = 0.0308, Welch’s ANOVA test with Dunnett’s T3 multiple-comparisons test; Fig. S3*B*), while REM duration and latency were restored relatively rapidly (REM duration: after stress (n = 4) vs 1W FLU (n = 4): 0.97 ± 0.10 vs 0.49 ± 0.09, *p* = 0.0293; 1W FLU (n = 4) vs 1M FLU (n = 4): 0.49 ± 0.09 vs 0.41 ± 0.11, *p* = 0.0233; *p* = 0.0064, Brown-Forsythe ANOVA test with Dunnett’s T3 multiple-comparisons test; Fig. S3*C*, REM latency: after stress (n = 4) vs 1W FLU (n = 4): 855.00 ± 327.83 vs 3427.50 ± 383.87, *p* = 0.0006; after stress (n = 4) vs 1M FLU (n = 4): 855.00 ± 327.83 vs 2617.50 ± 163.11, *p* = 0.0071; *p* = 0.0007, one-way ANOVA with Tukey’s multiple-comparisons test; Fig. S3*D*). Since these sleep alterations by SDS and recoveries by chronic AD treatments resembled those observed in patients with MDD [Benca et al., 1992; Benca, 2005; Peterson and Benca, 2008], these findings demonstrate the validity of the method as an MDD model from the standpoint of sleep architecture.

### EEG spectral power analyses

EEG spectral powers of REM sleep during the light phase (ZT0 – ZT8.5) were analyzed. SDS significantly decreased the long-range averaged relative REM sleep theta power (5 – 9 Hz) (control (n =5) vs stress (n =6): 0.038 ± 0.004 vs 0.027 ± 0.002, *p* < 0.0001; *F*_(45, 256)_ = 3.03, *p* < 0.0001, two-way ANOVA with Tukey’s multiple-comparisons test; Fig. 2*E, F*), while chronic AD treatment increased it (stress (n = 6) vs IMI (n = 5): 0.027 ± 0.002 vs 0.034 ± 0.003, *p* < 0.0001; *F*_(45, 256)_ = 3.03, *p* < 0.0001, two-way ANOVA with Tukey’s multiple-comparisons test; Fig. 2*E, F*). Furthermore, chronic FLU treatment increased the REM sleep theta powers in a time-dependent manner (stress (n =4) vs 1M FLU (n = 4): 0.030 ± 0.002 vs 0.035 ± 0.004, *p* = 0.0111; *F*_(15, 144)_ = 18.28, *p* < 0.0001, two-way ANOVA; Tukey’s multiple-comparisons test; Fig. S3*E, F*).

### Correlation between EEG sleep parameters and social interaction ratios

To evaluate which sleep parameters were responsible for the MDD-like social avoidance, we then performed correlation analyses between six sleep variables and social interaction ratios at 1M (Fig. 3*A*). We found that the interaction ratios, both to an unfamiliar BN or SD rat, at 1M after the last SDS, correlated significantly with the relative averaged REM sleep theta powers (BN: *r* = 0.6034, *p* = 0.0133; Fig. 3*B*, SD: *r* = 0.5731, *p* = 0.0203; Fig. 3*C*). No such significant correlation was found between the interaction ratios and the REM or NREM durations and bouts, or between the interaction ratios and the REM latency (Fig. 3*D*, S4*A, B*).

**Figure 3.**
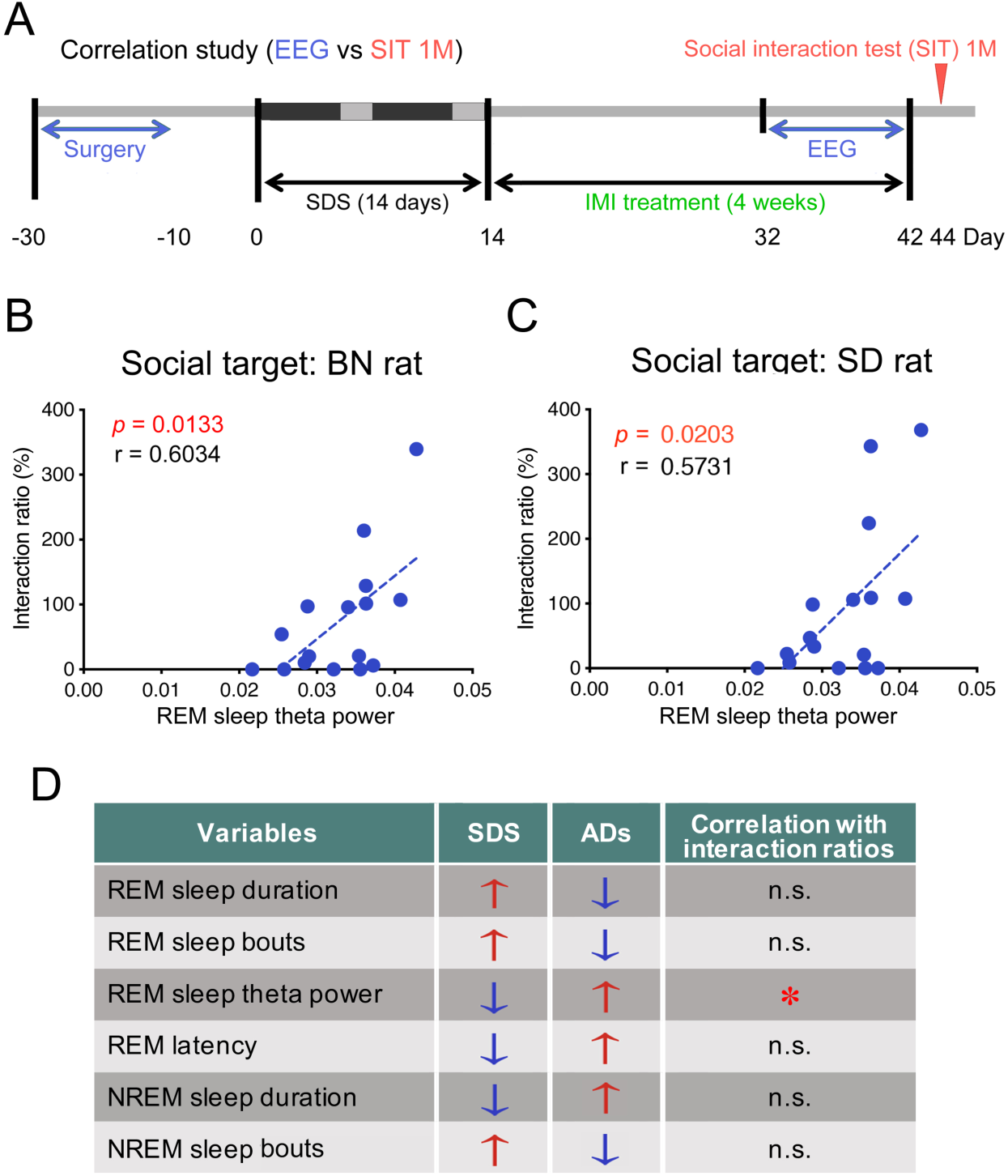
Correlations between REM sleep theta powers and interaction ratios one month after the last SDS. ***A***, Experimental schedule for correlation analysis. ***B, C***, Correlation between averaged relative REM theta powers during the light phase (ZT0 – ZT8.5) and interaction ratios against social targets (BN rats; ***B***, SD rat; ***C***) one month after the last SDS. ***D***, Summary of the correlation study (see Fig. S4). Note that, among various sleep variables, only the REM sleep theta power significantly correlated with social interaction ratios (*).

### Fear conditioning analysis

Recent evidence suggests that REM sleep theta oscillations might play a key role in the consolidation of negative/aversive memories [Hutchison and Rathore, 2015; Boyce et al., 2016]. We, therefore, examined the fear-conditioned contextual and cued memory formation one month after the last SDS (on Days 44 – 46; Fig. 4*A*). During conditioning (Day 1), freezing behavior between groups was not significantly different during conditioning in context A (between 120 – 300 sec: *F*_(2, 18)_ = 0.3172, *p* = 0.7322, one-way ANOVA; Fig. 4*B*) nor during the tones (tone 1; *F*_(2, 14)_ = 1.142, *p* = 0.3472, one-way ANOVA; Fig. 4*C*, and tone 2; *F*_(2, 14)_ = 1.254, *p* = 0.3155, one-way ANOVA; Fig. 4*D*). On Day 2, SDS significantly reduced freezing behavior (control (n =5) vs stress (n = 6): *p* = 0.0103; *F*_(2, 14)_ = 7.315, *p* = 0.0067, one-way ANOVA with Tukey’s multiple-comparisons test; Fig. 4*E*) in context A, suggesting that our SDS paradigm induced a contextual memory deficit even one month after the last SDS. Chronic IMI treatment significantly rescued this deficit (stress (n = 6) vs IMI (n =6): *p* = 0.0201; one-way ANOVA with Tukey’s multiple-comparisons test; Fig. 1*E*). On Day 3, rats were placed in a novel context (context B) for a total of 5 min for cued recall testing. Freezing behavior during the tone (2 min) did not differ between the two groups, with each showing a robust and selective freezing response based on the cue (tone) (control (n =5) vs stress (n = 6): *p* = 0.5347; *F*_(2, 14)_ = 0.9034, *p* = 0.4276, one-way ANOVA with Tukey’s multiple-comparisons test; Fig. 4*F*).

**Figure 4.**
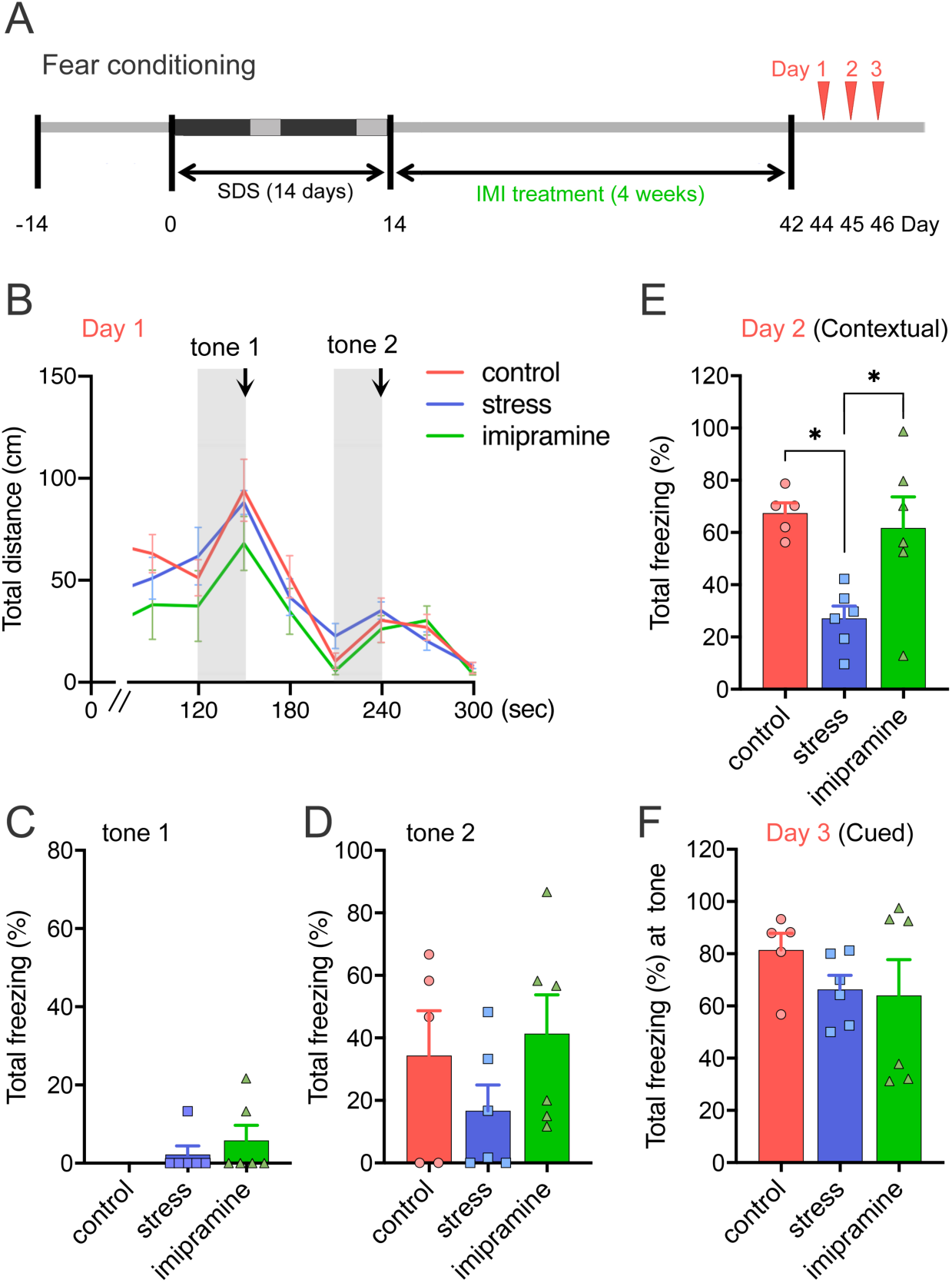
Memory function in the novel SDS rat model. ***A***, Experimental schedules for the fear conditioning test. ***B, C, D***, Fear conditioning test at one month after the last SDS (Days 44 – 46; control: n = 5; stress: n = 6; imipramine (IMI): n = 6). Arrows indicate a timing of 2 s of 0.3 mA current delivery. On Day 1, no group difference was found in the total distance moved (***B***) and the total freezing during tones 1 (***C***) and 2 (***D***). ***E***, Contextual recall memory test on Day 2. Recall impairment was found in the stressed rats, which was recovered by IMI. \**p* < 0.05, One-way ANOVA followed by Tukey’s test. ***F***, Cued recall memory test on Day 3.

## Discussion

The biological research for disturbed sleep in MDD has been largely fruitless despite almost 50 years of investigations. Inconsistent findings regarding the sleep disturbances in MDD and the effects of ADs on sleep, as well as misconceptualizations of EEG variables and animal models, may be fundamental barriers to the identification of true targets. This study represents an attempt to reconcile these inconsistencies and misconceptualizations by using an animal model. Specifically, our data point to the possibility that, among various sleep parameters, REM sleep theta power, a parameter never measured directly in MDD patients, strongly correlates with the MDD-like behaviors we have observed in rats.

In the present study, we applied the 14-day SDS paradigm to rats, focusing on the post-stress effects, and found that SDS induces long-lasting behavioral abnormalities in association with MDD-like sleep alterations even a month after the end of SDS, which are largely restored by chronic treatments with two different classes of ADs, imipramine and fluoxetine. In addition, our model has several unique characteristics resembling human MDD. First, although chronic AD treatments significantly improved SDS-induced social avoidance, about 30-40% of non-responders were always observed in the social interaction tests (Fig. 1*B, C*). Second, the pattern of disturbed sleep in our model closely resembles those found in MDD. Third, ADs restored most of the MDD-like sleep alterations except for the shortened REM sleep latency (Fig. 2*D*). Longitudinal studies have shown shortened REM sleep latency to be stable within depressed individuals over time, regardless of clinical state [Rush et al., 1986]. Taken together, we concluded that this rat model fulfills the face, construct, and predictive validities of a model for disturbed sleep in MDD.

Notably, among various sleep parameters, REM sleep theta powers most significantly correlated with the social interaction ratios (Fig. 4*B, C*), suggesting a possible role of REM sleep theta oscillation in mood regulation. A pioneering study by Hegde et al. investigated the effects of 2-hr chronic immobilization stress (CIS) for 10 days on REM sleep theta oscillation and found that a subgroup of stressed rats showed increased REM sleep durations in association with attenuated REM sleep theta powers [Hegde et al., 2011]. Recently, four independent studies, using mouse SDS [Wells et al., 2017; Henderson et al., 2017], unpredictable chronic mild stress (UCMS) [Nollet et al., 2019], and water immersion and restraint stress (WIRS) [Yasugaki et al., 2019] models, have shown that these stresses induced changes in various REM sleep parameters. These changes, however, were observed mostly during the stress periods and little was observed in terms of post-stress effects (only 5 days after the last SDS [Wells et al., 2017; Henderson et al., 2017]). Furthermore, none of these studies reported the effects of chronic AD treatments.

The effects of various stresses on REM sleep parameters also proved inconclusive in these studies. In the first mouse SDS model [Wells et al., 2017], REM sleep duration, bouts, and theta power (per day) all increased during SDS while REM sleep theta power appeared to decrease on day 10 of the SDS. In the other mouse SDS model [Henderson et al., 2017], REM sleep durations and bouts decreased and REM sleep theta power remained unchanged during the light phase. In the UCMS model [Nollet et al., 2019], REM sleep duration, bouts, and theta power (per day) increased during the 9-week UCMS. In the WIRS model [Yasugaki et al., 2019], REM sleep duration and bouts during the dark phase increased during the first week of the stress episode. REM sleep theta power, during both the dark and light phases, decreased and this reduction was more pronounced during the second and third weeks of the stresses compared with that during the first week [Yasugaki et al., 2019].

In our rat SDS model, REM sleep duration and bouts increased while REM sleep theta power decreased during the light phase for up to 1M post-SDS period, which were all recovered by chronic AD (IMI & FLU) treatments. One possible explanation for these differences between ours and the above-mentioned studies may be the use of rats plus the 14-day SDS procedure. While rats have a clearly defined hierarchy system that finds its expression through submissive postures or attacks, mice do not typically present such hierarchical behaviors [Dixon, 2004].

We further found that chronic FLU treatment gradually increased the REM sleep theta power in a time-dependent manner (Fig. S3*E, F*), which contrasted strikingly to the relatively rapid (1 week) effects of FLU on the REM duration (Fig. S3*C*) and REM latency (Fig. S3*D*). This suggests that slower changes take place within the brain areas regulating REM sleep theta power (e.g., basal forebrain and hippocampus) compared to those regulating REM sleep timing (e.g., brainstem, midbrain, and hypothalamus). In the basal forebrain, GABAergic neurons in the medial septum (MS)/diagonal band of the Broca are believed to play critical roles in the generation of hippocampal REM theta oscillations [Lawson and Bland, 1993; Dragoi et al., 1999; Simon et al., 2006; Hangya et al., 2009; Zhang et al., 2011]. More recently, Boyce et al. have demonstrated that the optogenetic silencing MS GABAergic neurons during REM sleep reduced REM sleep theta power in the hippocampus, resulting in the subsequent erasure of novel object place recognition and impaired fear-conditioned contextual (but not cued) memory [Boyce et al., 2016]. Consistently, we also found similar contextual (but not cued) fear memory deficits in the SDS rats at 1M, which was rescued by IMI (Fig. 4), in association with the REM sleep theta power changes (Fig. 2*F*). These findings give rise to the possibility that the septohippocampal pathway [Manseau et al., 2008; Grosmark et al., 2012] may be largely impaired by SDS and restored gradually by chronic AD treatments.

Finally, Montgomery et al. [2008] have reported that the subregional theta-gamma EEG pattern in REM sleep, compared with waking exploration, suggests enhanced offline processing in the dentate gyrus (DG) and CA3 regions of the hippocampus during REM sleep. Since chronic stress reduces DG neurogenesis and CA3 dendritic arborization while chronic AD treatments restore them [Anacker and Hen, 2017; McEwen, 2017], this type of intrinsic intrahippocampal theta synchronization may also be involved in the changes in the REM sleep theta power in our model.

Limitations of the current study should be noted. First, although we have tentatively concluded, based on the patterns of disturbed sleep and the effects of ADs, that ours is an effective model for MDD, it may still better serve as a model for other psychiatric disorders such as post-traumatic stress disorders. Second, we did not include experimental groups (Control + ADs) in the current study. This is partly because enormous numbers of such studies have been reported since the 1950s (e.g., Gao et al., 1992) and the only likely conclusion is that ADs suppress REM sleep time. Third, all the data in the current study is correlational but not causal in nature.

In conclusion, we have established a novel SDS rat model for disturbed sleep in MDD and have provided validatory evidence from the standpoint not only of behaviors, but also of sleep architectures. Our EEG analyses demonstrate that chronic AD treatments ameliorated MDD-like sleep disturbances. Furthermore, ADs restored MDD-like social avoidance, potentially by improving the REM sleep theta power. Although further validation in humans is required, our model will be useful for the development of new pharmacotherapies for disturbed sleep in MDD.

## Acknowledgments

We thank Prof. Adel C. Alonso for her critical reading. We also thank Ms Moe Watanabe for her technical assistance. This work was funded by JSPS KAKENHI Grant Numbers 25871162 and 16K04442 for Y.M. and 15K15438 for Y.T.

## Author contributions

Y.M. and Y.T. designed the experiments; Y.M., N.O., T.S., K.A., N.N., and Y.T. performed the experiments; Y.M., N.O, T.S, and Y.T. designed the analyses and discussed the results; Y.M. and Y.T. analyzed the data; Y.T. wrote the paper; Y.M. and N.O. performed animal surgeries; and all authors commented on the manuscript.

## References

Azmitia EG, Siegel M (1978) An autoradiographic analysis of the differential ascending projection of dorsal and medial raphe nuclei in the rat. Comp Neurol 179:641–668.

Baglioni C, Nanovska S, Regen W, Spiegelhalder K, Feige B, Nissen C, Reynolds CF, Riemann D (2016) Sleep and mental disorders: A meta-analysis of polysomnographic research. Psychol Bul 142:969–990.

Beitinger ME, Fulda S (2010). Long-term effects of antidepressants on sleep. In: Sleep and Mental Illness: Pandi-Perumal SR, Kramer M (ed), pp 183–201. New York: Cambridge University Press.

Benca RM (2005) Mood disorders. In: Principles and practice of sleep medicine. 4th ed: Kryger MH, Roth T, Dement WC (ed), pp1311–1326. Philadelphia: Elsevier Saunders

Benca RM, Obermeyer WH, Thisted RA, Gillin JC (1992) Sleep and psychiatric disorders: a meta-analysis. Arch Gen Psychiatry 49:651–68.

Berton O, McClung CA, Dileone RJ, Krishnan V, Renthal W, Russo SJ, Graham D, Tsankova NM, Bolanos CA, Rios M, Monteggia LM, Self DW, Nestler EJ (2006) Essential role of BDNF in the mesolimbic dopamine pathway in social defeat stress. Science 311:864–868.

Boyce R, Glasgow SD, Williams S, Adamantidis A (2016) Causal evidence for the role of REM sleep theta rhythm in contextual memory consolidation. Science 352:812–816.

Cao JL, Covington HE, Friedman AK, Wilkinson MB, Walsh JJ, Cooper DC, Nestler EJ, Han MH (2010) Mesolimbic dopamine neurons in the brain reward circuit mediate susceptibility to social defeat and antidepressant action. J Neurosci 30:16453–16458.

Der-Avakian A, Mazei-Robison MS, Kesby JP, Nestler EJ, Markou A (2014) Enduring deficits in brain reward function after chronic social defeat in rats: susceptibility, resilience, and antidepressant response. Biol Psychiatry 76:542–549.

Dixon AK (2004) The Social Behaviour of Mice and its Sensory Control. In: H Hedrich (ed). The laboratory mouse, pp. 287–300. Academic Press: London.

Dragoi G, Carpi D, Recce M, Csicsvari J, Buzsáki G (1999) Interactions between hippocampus and medial septum during sharp waves and theta oscillation in the behaving rat. J Neurosci 19:6191–6199.

Ford DE, Kamerow DB (1989) Epidemiologic study of sleep disturbances and psychiatric disorders: An opportunity for prevention? JAMA 262:1479–1484.

Gao B, Duncan WC Jr, Wehr TA (1992) Fluoxetine decreases brain temperature and REM sleep in Syrian hamsters. Psychopharmacology (Berl) 106:321–329.

Grosmark AD, Mizuseki K, Pastalkova E, Diba K, Buzsáki G (2012) REM sleep reorganizes hippocampal excitability. Neuron 75:1001–1007.

Hangya B, Borhegyi Z, Szilágyi N, Freund TF, Varga V (2009) GABAergic neurons of the medial septum lead the hippocampal network during theta activity. J Neurosci 29:8094–8102.

Harper DG, Tornatzky W, Miczek KA (1996) Stress induced disorganization of circadian and ultradian rhythms: comparisons of effects of surgery and social stress. Physiol Behav 59:409–419.

Hegde P, Jayakrishnan HR, Chattarji S, Kutty BM, Laxmi TR (2011) Chronic stress-induced changes in REM sleep on theta oscillations in the rat hippocampus and amygdala. Brain Res 1382:155–164.

Henderson F, Vialou V, Mestikawy SE, Fabre V (2017) Effects of social defeat stress on sleep in mice. Front Behav Neurosci 11:227.

Hutchison IC, Rathore S (2015) The role of REM sleep theta activity in emotional memory. Front Psychol 6: 1439.

Jones BE (1991) Paradoxical sleep and its chemical/structural substrates in the brain. Neuroscience 40:637–656.

Krishnan V, Han MH, Graham DL, Berton O, Renthal W, Russo SJ, Laplant Q, Graham A, Lutter M, Lagace DC, Ghose S, Reister R, Tannous P, Green TA, Neve RL, Chakravarty S, Kumar A, Eisch AJ, Self DW, Lee FS, Tamminga CA, Cooper DC, Gershenfeld HK, Nestler EJ (2007) Molecular adaptations underlying susceptibility and resistance to social defeat in brain reward regions. Cell 131:391–404.

Krystal AD, Thakur M, Roth T (2008) Sleep disturbance in psychiatric disorders: effects on function and quality of life in mood disorders, alcoholism, and schizophrenia. Ann Clin Psychiatry 20:39–46.

Kupfer DJ, Shaw DH, Ulrich R, Coble PA, Spiker DG (1982) Application of automated REM analysis in depression. Arch Gen Psychiatry 39:569–573.

Landolt HP, Kelsoe JR, Rapaport MH, Gillin JC (2003) Rapid tryptophan depletion reverses phenelzine-induced suppression of REM sleep. J Sleep Res 12:13–18.

Lawson VH, Bland BH (1993) The role of the septohippocampal pathway in the regulation of hippocampal field activity and behavior: Analysis by the intraseptal microinfusion of carbachol, atropine, and procaine. Exp Neurol 120:132–144.

McEwen BS (2017) Neurobiological and systemic effects of chronic stress. Chronic Stress 1:1–11.

Manseau F, Goutagny R, Danik M, Williams SJ (2008) The hippocamposeptal pathway generates rhythmic firing of GABAergic neurons in the medial septum and diagonal bands: an investigation using a complete septohippocampal preparation in vitro. Neurosci 28:4096–4107.

Mayers AG, Baldwin DS (2005) Antidepressants and their effect on sleep. Hum Psychopharmacol 20:533–539.

McGinty DJ, Harper RM (1976) Dorsal raphe neurons: depression of firing during sleep in cats. Brain Res 101:569–575.

Meerlo P, Sgoifo A, Turek FW (2002) The effects of social defeat and other stressors on the expression of circadian rhythms. Stress 5:15–22.

Meerlo P, Van den Hoofdakker RH, Koolhaas JM, Daan S (1997) Stress-induced changes in circadian rhythms of body temperature and activity in rats are not caused by pacemaker changes. J Biol Rhythms 12:80–92.

Montgomery SM, Sirota A, Buzsáki G (2008) Theta and gamma coordination of hippocampal networks during waking and rapid eye movement sleep. J Neurosci 28:6731–6741.

Moussavi S, Chatterji S, Verdes E, Tandon A, Patel V, Ustun B (2007) Depression, chronic diseases, and decrements in health: results from the world health surveys. Lancet 370:851–858.

Nollet M, Hicks H, McCarthy AP, Wu H, Möller-Levet CS, Laing EE, Malki K, Lawless N, Wafford KA, Dijk DJ, Winsky-Sommerer R (2019) REM sleep’s unique associations with corticosterone regulation, apoptotic pathways, and behavior in chronic stress in mice. Proc Natl Acad Sci USA 116:2733–2742.

Oikonomou G, Altermatt M, Zhang RW, Coughlin GM, Montz C, Gradinaru V (2019) The Serotonergic Raphe Promote Sleep in Zebrafish and Mice. Neuron 103:686–701.

Palagini L, Baglioni C, Ciapparelli A, Gemignani A, Riemann D (2013) REM sleep dysregulation in depression: State of the art. Sleep Med Rev 17:377–390.

Peterson MJ, Benca RM (2008) Sleep in mood disorders. Sleep Med Clin 3:231–249.

Pillai V, Kalmbach DA, Ciesla JA (2011) A meta-analysis of electroencephalographic sleep in depression: Evidence for genetic biomarkers. Biol Psychiatry 70:912–919.

Riemann D, Berger M, Voderholzer U (2001) Sleep and depression - results from psychobiological studies: an overview. Biol Psychol 57:67–103.

Rush AJ, Erman MK, Giles DE, Schlesser MA, Carpenter G, Vasavada N, Roffwarg HP (1986) Polysomnographic findings in recently drug-free and clinically remitted depressed patients. Arch Gen Psychiatry 43:878–884.

Sakai K (2011) Sleep-waking discharge profiles of dorsal raphe nucleus neurons in mice. Neuroscience 197:200–224.

Scammell TE, Arrigoni E, Lipton JO (2017) Neural circuitry of wakefulness and sleep. Neuron 93: 747–765.

Shinba T (2009) 24-h profiles of direct current brain potential fluctuation in rats. Neurosci Lett 465:104–107.

Simon AP, Poindessous-Jazat F, Dutar P, Epelbaum J, Bassant MH (2006) Firing properties of anatomically identified neurons in the medial septum of anesthetized and unanesthetized restrained rats. J Neurosci 26:9038–9046.

Stassen HH, Delini-Stula A, Angst J (1993) Time course of improvement under antidepressant treatment: a survival-analytical approach. Eur Neuropsychopharmacol 3:127–135.

Thakkar MM, Strecker RE, McCarley RW (1998) Behavioral state control through differential serotonergic inhibition in the mesopontine cholinergic nuclei: a simultaneous unit recording and microdialysis study. J Neurosci 18:5490–5497.

Thase ME, Buysse DJ, Frank E, Cherry CR, Cornes CL, Mallinger AG, Kupfer DJ (1997) Which depressed patients will respond to interpersonal psychotherapy? The role of abnormal EEG sleep profiles. Am J Psychiatry 154:502–509.

Tornatzky W, Miczek KA (1993) Long-term impairment of autonomic circadian rhythms after brief intermittent social stress. Physiol Behav 53:983–993.

Tsankova NM, Berton O, Renthal W, Kumar A, Neve RL, Nestler EJ (2006) Sustained hippocampal chromatin regulation in a mouse model of depression and antidepressant action. Nat Neurosci 9:519–525.

Tsuno N, Besset A, Ritchie K (2005) Sleep and depression. J Clin Psychiatry 66:1254–1269.

Vialou V, Robison AJ, LaPlant QC, Covington HE, Dietz DM, Ohnishi YN, Mouzon E, Rush AJ 3rd, Watts EL, Wallace DL, Iñiguez SD, Ohnishi YH, Steiner MA, Warren BL, Krishnan V, Bolaños CA, Neve RL, Ghose S, Berton O, Tamminga CA, Nestler EJ (2010) ΔFosB in brain reward circuits mediates resilience to stress and antidepressant responses. Nat Neurosci 13:745–752.

Weissbourd B, Ren J, DeLoach KE, Guenthner KJ, Miyamichi K, Luo L (2014) Presynaptic Partners of Dorsal Raphe Serotonergic and GABAergic Neurons. Neuron 83:645–662.

Wells AM, Ridener E, Bourbonais CA, Kim W, Pantazopoulos H, Carroll FI, Kim KS, Cohen BM, Carlezon WA Jr (2017) Effects of chronic social defeat stress on sleep and circadian rhythms are mitigated by kappa-opioid receptor antagonism. J Neurosci 37:7656–7668.

Wilkinson MB, Xiao G, Kumar A, LaPlant Q, Renthal W, Sikder D, Kodadek TJ, Nestler EJ (2009) Imipramine treatment and resiliency exhibit similar chromatin regulation in the mouse nucleus accumbens in depression models. J Neurosci 29:7820–7832.

Yasugaki S, Liu CY, Kashiwagi M, Kanuka M, Honda T, Miyata S, Yanagisawa M, Hayashi Y (2019) Effects of 3 weeks of water immersion and restraint stress on sleep in mice. Front Neurosci 13:1072.

Zhang H, Lin SC, Nicolelis MA (2011) A distinctive subpopulation of medial septal slow-firing neurons promote hippocampal activation and theta oscillations. J Neurophysiol 106:2749–2763.

